# Optimal protocols for sequence-based characterization of the human vaginal microbiome

**DOI:** 10.1101/2020.05.05.079996

**Authors:** Luisa W. Hugerth, Marcela Pereira, Yinghua Zha, Maike Seifert, Vilde Kaldhusdal, Fredrik Boulund, Maria C Krog, Zahra Bashir, Marica Hamsten, Emma Fransson, Henriette Svarre-Nielsen, Ina Schuppe-Koistinen, Lars Engstrand

**Affiliations:** Centre for Translational Microbiome Research (CTMR), Department of Microbiology, Tumour and Cell Biology (MTC), Karolinska Institutet, Science for Life Laboratory, Solna, Sweden; Department of Medicine Solna, Division of Infectious Diseases, Karolinska University Hospital, Center for Molecular Medicine, Karolinska Institutet, Solna, Sweden; The Recurrent Pregnancy Loss Unit, Capital Region of Denmark, Rigshospitalet and Hvidovre Hospital, Copenhagen, Denmark; Department of Clinical Immunology, Copenhagen University Hospital, Denmark; Department of Obstetrics and Gynaecology, Holbæk Hospital, Holbæk, Denmark; Department of Women’s and Children’s Health, Uppsala university, Uppsala, Sweden; Department of Obstetrics and Gynecology, Hvidovre Hospital, Copenhagen Denmark

## Abstract

The vaginal microbiome has been connected to a wide range of health outcomes. This has led to a thriving research environment, but also to the use of conflicting methodologies to study its microbial composition. Here we systematically assess best practices for the sequencing-based characterization of the human vaginal microbiome. As far as 16S rRNA gene sequencing is concerned, the V1-V3 region has the best theoretical properties, but limitations of current sequencing technologies mean that the V3-V4 region performs equally well. Both of these approaches present very good agreement with qPCR quantification of key taxa, provided an appropriate bioinformatic pipeline is used. Shotgun metagenomic sequencing presents an interesting alternative to 16S amplification and sequencing, but it is not without its challenges. We have assessed different tools for the removal of host reads and the taxonomic annotation of metagenomic reads, including a new, easy-to-build and – use, reference database of vaginal taxa. This strategy performed as well as the best performing previously published strategies. Despite the many advantages of shotgun sequencing none of the shotgun approaches assessed here had as good agreement with the qPCR data as 16S rRNA gene sequencing.

**Importance:** The vaginal microbiome has been connected to a wide range of health outcomes, from susceptibility to sexually transmitted infections to gynecological cancers and pregnancy outcomes. This has led to a thriving research environment, but also to conflicting available methodologies, including many studies that do not report their molecular biological and bioinformatic methods in sufficient detail for them to be considered reproducible. This can lead to conflicting messages and delay progress from descriptive to intervention studies. By systematically assessing best practices for the characterization of the human vaginal microbiome, this study will enable past studies to be assessed more critically and assist future studies in the selection of appropriate methods for their specific research questions.

## Introduction

The human vaginal microbiome plays a key role in maintaining the gynecological health of in women of reproductive age. Estrogen is responsible for the cyclic maturation of the vaginal epithelium and the deposition of glycogen in vaginal epithelial cells (1). Shedded glycogen-rich cells are an excellent carbon source for lactic acid bacteria (2). Lactic acid lowers the local pH and has bactericidal and immune regulatory effects (3). In addition to keeping bacterial balance and preventing bacterial vaginosis (BV) and aerobic vaginitis (AV) (4), the vaginal microbiome has been shown to play a protective role against viral infections such as HPV (5), HSV-2 (6) and HIV (7). It might also be protective against adverse pregnancy outcomes such as early miscarriage (8) and preterm birth (9) as well as gynecological cancers (10).

In clinical practice, the diagnosis of bacterial vaginosis is often based on experienced vaginal symptoms and pH-testing, sometimes combined with a visual assessment of a vaginal smear wet mount under microscopy. Criteria such as the Amsel criteria (11) and Nugent scoring (12) have been developed to assist in this assessment, but are low resolution and low throughput. In research settings, for the past 10 years it has become standard to sequence part of the 16S rRNA gene to characterize the vaginal microbiome in high throughput. However, no consensus exists in this field for experimental or bioinformatic best practices, with different studies (sometimes within the same research group) focusing on different variable regions of the 16S rRNA gene (see **Table 1**) (13–20).

While extensive work has been published assessing best practices for characterizing free living bacterial communities (21) or human-associated microbes as a whole (15), these findings are not directly translatable to the human vaginal microbiome for a few reasons. Firstly, clinically important species such as Mycoplasma genitalium or Chlamydia trachomatis have an unusual pattern of substitutions in their rRNA gene, meaning that optimizing for a broad taxonomic range might have the unwanted effect of missing these species. Even more importantly, the 16S rRNA gene is generally regarded to only provide taxonomic resolution down to the genus level (22). However, for the human vaginal microbiome, distinguishing between different Lactobacillus species is crucial, since eg Lactobacillus crispatus often plays a protective role not exerted by Lactobacillus iners (5, 7, 23).

One way to bypass the trade-offs involved in selecting a PCR primer set is to perform full metagenomic shotgun sequencing. This approach presents several advantages and some serious challenges. Among the advantages of metagenomics is the possibility of going deeper than species-level classification, including the identification of strains and specific genes. Indeed, recent work applying metagenomics to a large set of vaginal samples has identified extensive intra-species variation in several important taxa such as various *Lactobacillus* species, *Gardnerella vaginalis* and *Atopobium vaginae* (24). It is also known that the degree of stability of the vaginal microbiome can be quite different between individuals (25), and subspecies resolution might be needed to explain why certain microbiomes are more resilient than others.

While all of the methods described above have the capability of broadly assessing a wide range of species, they are only semiquantitative and may introduce different biases at the library preparation and bioinformatics steps. To systematically assess the effect of different variable regions, different bioinformatic approaches and different taxonomic annotation pipelines on the observed microbial profile of human vaginal samples, we have attempted to identify all primer pairs used in published human vaginal microbiome studies in the past decade. Each of these primers was assessed *in silico* for taxonomic coverage and annotation accuracy. Different annotation schemes were used for each primer pair. The pairs with the best performance were taken into the lab and used to amplify the same set of samples. Furthermore, shotgun metagenomic sequencing was applied to each of these sample as well. This way, we can directly compare the results between primer sets and sequencing strategies..

The gold standard for quantifying specific organisms is still qPCR, a fully quantitative method. Here we have performed qPCR on three key vaginal taxa (*Lactobacillus crispatus, Lactobacillus iners* and *Gardnerella vaginalis*), to provide a ground truth against which each of the other methods can be assessed. The results described here can guide the implementation of future vaginal microbiome studies, as well as providing important information for the comparison of previous studies which have used diverging methods.

## Material and Methods

### Construction of the databases

To create a corresponding shotgun database, we started from the list of vaginotropic species published by Diop *et al* (26). In addition to these previously published results, a dataset of 480 vaginal swabs collected throughout the menstrual cycle of a healthy Danish cohort (Elkjær-Madsen *et al*, submitted) sequenced by CoreBiome (St Paul, MN, USA) using BoosterShot technology was used. For every bacterial species identified in the dataset and not present in the Diop database, manual searches of PubMed and NucCore were made, and the species was kept if it had been previously identified in the human urogenital tract. Eukaryotic species were added by searching NucCore with the search key “((vagina[All Fields] AND “Eukaryota”[Organism]) NOT “Metazoa”[Organism]) NOT “Viridiplantae”[Organism] AND (biomol_genomic[PROP] AND refseq[filter])”. Finally, a free-text search for “BVAB” retrieved metagenome-associated genomes representative of the bacterial-vaginosis associated Clostridiales group. The resulting list of taxa is available as **Suppl. Table 1**. When a taxon could not be programmatically included in the database, manual searches against NCBI’s Taxonomy database were used to verify whether the taxon name had been updated. Not all taxa could be retrieved as full genomes, as some are present in the databases only as single genes, and they are therefore missing from the current version of the database. The resulting database (v0.1) and the scripts used for producing a genome database based on a taxon list are available at https://github.com/ctmrbio/optivag/tree/master/database

### Simulated amplicons

Amplicons were extracted from the 16S rRNA gene database based on exact matches to the primers. For amplicons starting at the 27f position, which is often not included in the reference sequence due to its location, two alternative approaches were compared. The pessimistic approach extracts only sequences containing the primer regions, while the optimistic assumes that all sequences lacking the 5’-end would be amplified by the 27f primer. The truth is likely somewhere in the middle of these extremes.

Reads were simulated for each primer pair using a read length of 250 bp. While it is possible to sequence longer fragments with the commercial kits available today, this is a realistic read length after trimming primer pairs and low-quality base pairs. For amplicons <500 bp long, the resulting reads were merged; otherwise, they were treated independently. When the resulting amplicon length was very close to 500 bp, both approaches were considered, since the ability to merge them becomes dependent on the accuracy of the sequencer used.

### Sample collection

Women were recruited by advertisements in student magazines, university notice boards and social media and included between September 2017 and January 2018 at Rigshospitalet, Copenhagen, Denmark. The women were provided with self-collection kits and received the needed instructions for the vaginal swab collection. In short, they were instructed to separate the labia major with one hand (in order to reduce the risk of contamination with microbiota from external genitals), insert a swab (FLOQSwabs, CP520CS01 Copan Flock Technologies, Brescia, Italy) in the vagina with the other hand and rotate it for 10-15 seconds before placing the swab into the provided collection tube (FluidX tubes; 65-7534 - Brooks Life Sciences, Chelmsford, MA, USA; containing 0.8 ml DNA/RNA-shield; R1100-250, Zymo Research, Irvine, CA, USA) and breaking off the handle. Samples were kept in room temperature for up to 2 weeks and then at -20°C for up to 4 weeks, before being transferred to -80°C. All participants gave oral and written consent to participate in the study and were remunerated with 3,000 DKK after completing sample collection. All data were collected and managed using REDCap electronic data capture tools (27), hosted at the Capital Region of Denmark. The study is approved by The Regional Committee on Health Research Ethics (H-17017580) and the Data Protection Agency in the Capital Region of Denmark (2012-58-0004).

### DNA extraction

DNA extraction was performed with the Quick-DNA Magbead Plus kit (D4082 - Zymo Research, Irvine, CA, USA), according to manufacturer instructions with few modifications. Prior to extraction the samples were bead-beaten for 1 minute at 1600 RPM using ZR Bashing Bead Lysis matrix (S6012 - Zymo Research, Irvine, CA, USA). After bead beating, samples were treated with a lysozyme solution 37°C for 60 min (lysozyme recipe: 20 mM Tris-CL, pH8; 2 mM sodium EDTA (Tris-EDTA - T9285); lysozyme (L6876-100G) to 100 mg/ml) and proteinase K at 55°C for 30 min (20 mg/ml, part of extraction kit), previously to DNA clean-up using a Freedom EVO robot (Tecan, Männendorf, Switzerland). Eight sample pools were created for this manuscript, consisting of 4 consecutive daily vaginal swabs from each of 8 individuals from a cohort of healthy young women. All eight sample pools were used for each of the experimental approaches attempted.

### Sequence amplification, sequencing and error correction

The following PCR set-ups were used:

- One-step PCR amplification of the V3-V4 region
- Two-step PCR amplification of the V3-V4 region
- Two-step PCR amplification of the V1-V3 region using reverse primer 515r
- Two-step PCR amplification of the V1-V3 region using reverse primer 534r
- The same settings as above were used for an experiment with a Chlamydia DNA spike-in (gblocks Gene Fragment, Integrated DNA Technologies (IDT), Coralville, IA, USA). DNA was spiked in at 1%, 5% or 10%.

The primer sequences and specific PCR conditions are described in **Supplementary Table 2.** All PCR reactions were perfomed with in 50 µL reactions using Phusion Hot Start II High-Fidelity PCR Master Mix (F-565L, Thermo Fisher Scientific, MA, USA). The 1-step PCR included 1.5 µL of DMSO. All PCR products were purified with Agencourt AMPure XP beads (A63881 - Beckman Coulter, Brea, CA, USA). For the two-step reactions, the purified sample was used as the template for barcoding with Nextera XT Index Kit v2 (FC-131-1002 - Illumina Inc, San Diego, CA, USA). The finished libraries were normalized to 4 nM and pooled, and were sequenced in a MiSeq using V4 chemistry (Illumina Inc).

Cutadapt (28) was used to trim primers, remove sequences not containing the expected primer pairs and remove bases with a Phred score <15. Error correction was performed with DADA2 (29) or Unoise (30), as described in the results.

### Taxonomic annotation of amplicons

Taxonomic annotation was performed with DADA2’s (v1.5) (29) built-in sequence classifier, based on the SILVA database (v128) (31) *In silico* amplicons were also classified by direct mapping to the SILVA v128 database.

For amplicons that could not be merged, a consensus between the potentially distinct annotations of forward and reverse reads was established as follows:

- If the two annotations were incompatible, the lowest common ancestor was kept (eg in case families agree, but genera diverge, only family-level annotation was kept)
- If one annotation was more detailed than the other (eg to genus level vs. to family level), but the two annotations agreed on all levels where they overlapped, the most detailed annotation was kept
- For species-level annotations where more than one species was possible, the intersection of the species suggested for each of the reads was kept (eg if the forward read is annotated as “Lactobacillus crispatus/gasseri/jenseni” and the reverse as “Lactobacillus gasseri/jenseni/longum”, the resulting annotation would be “Lactobacillus gasseri/jenseni”)

### Metagenomic shotgun sequencing

The same eight pools were used for whole genome library preparation. MGI FS DNA library prep kit (16x, 1000006987; MGI, Shenzhen - China) was used according to manufacturer instructions, except that 50 ng of DNA were used as input instead of the suggested 200 ng. Due to the smaller amount of input DNA, instead of double bead clean-up for size selection, a single clean-up step is applied. This kit uses enzymatic fragmentation of DNA followed by barcoding of samples (using 7 PCR cycles), single strand circularization and DNA nanoball construction. All procedures were automated using SP-960 and SP-100 robots (MGI). The sequencing step was performed in a DNBSEQ-G400 sequencer (MGI) using the high-throughput sequencing set (PE150 1000016952; MGI) with DNA libraries loaded onto to the flow cell using the DNB-loader MGIDL-200 (MGI).

### Human DNA removal

Human reads were removed *in silico* by one of the following strategies:

- BMTagger v1.1.0 (32) mapping to the GRCh38 reference library with standard masking
- BBMap v38.68 (33) against the hg19 reference library, masked as described in http://seqanswers.com/forums/showthread.php?t=42552
- Kraken2 v2.0.8-beta (34) against its built-in GRCh38 human reference, setting the confidence parameter to 0.1
- Kraken2 with the same parameters as above, but adding flag --quick

To independently be able to assess the human read removal performance of the aforementioned methods, reads were mapped to the hg19 masked reference using Bowtie2 v2.3.5 (35).

### Taxonomic annotation of shotgun reads

For assigning taxonomy to the remaining microbial reads, four approaches were assessed:

- Metaphlan2 v2.9.21 (36) with standard parameters
- Kraken2 v2.0.8-beta (34) to a general database (built using –download-library flags for archaea, bacteria, viruses, fungi and human) setting confidence to 0.5, followed by Bracken v2.0 (37) with threshold set to 1 read per million
- Kraken2 with the same parameters, but using the curated vaginal database described above
- Metalign v0.9.1 (38) with length normalization

### qPCR quantification of key taxa

To further validate the results observed by sequencing, three key taxa, namely *Lactobacillus crispatus* (VPI-3199), *Lactobacillus iners* (ATCC-55195) and *Gardnerella vaginalis* (CCUG-44120) were quantified by qPCR using LightCycler 480 (Roche, Mannheim – Germany) and SYBR Green assay from Bio-Rad (1725270 - BioRad, Sundbyberg - Sweden). The primer sequences and PCR conditions are described in **Supplementary Table 2**. In the triax-plots presented, the sum of these three taxa is normalized to 1 for each method presented, to allow a direct comparison.

## Results and discussion

### Coverage of each primer

To assess how well each primer sequence or primer combination covers potential vaginal taxa, all sequences matching each primer or primer combination were extracted from the database with a regular expression. A problem for the 27f primer variants is that many sequences in the database are incomplete at their 5’-end, which makes this assessment impossible. The same was not true at the 3’-end: the coverage for this region does not wane until after the V8 region, so it did not affect the assessment of any primers. The total coverage of each individual primer is depicted in **table 2**, and coverage for primer pairs in **table 3**. It is clear that pair 967-1061 performs much poorer than the remainder, with the exception of the 27f primers which could not be properly assessed.

In addition to covering a large percentage of all sequences, it is important that primers avoid taxonomic bias. The taxonomic coverage of each primer pair is depicted in **suppl. table 3**. Three of the genera that are mostly missed are Propionibacterium, Chlamydia and Mycoplasma. Propionibacteria are well covered by 341f-805r and possibly 27f-338r. These same pairs perform well with Mycoplasma, but only the former also covers Chlamydia. To add Chlamydia coverage to the 27f pool, one extra degeneracy has to be added to the reverse primer, making it either 515r 6×5’-GTGBCAGCMGCCGCGGTAA-3’ + 5’-GTGCCAGCAGCTGCGGTAA-3’ or 534r 5’-GTGCCAGCAGCYGCGGTAA-3’. **Fig. 1** shows a heatmap with the taxonomic coverage of each primer pair, assuming match of the 27f primers, where assessment was impossible.

**Figure 1.**
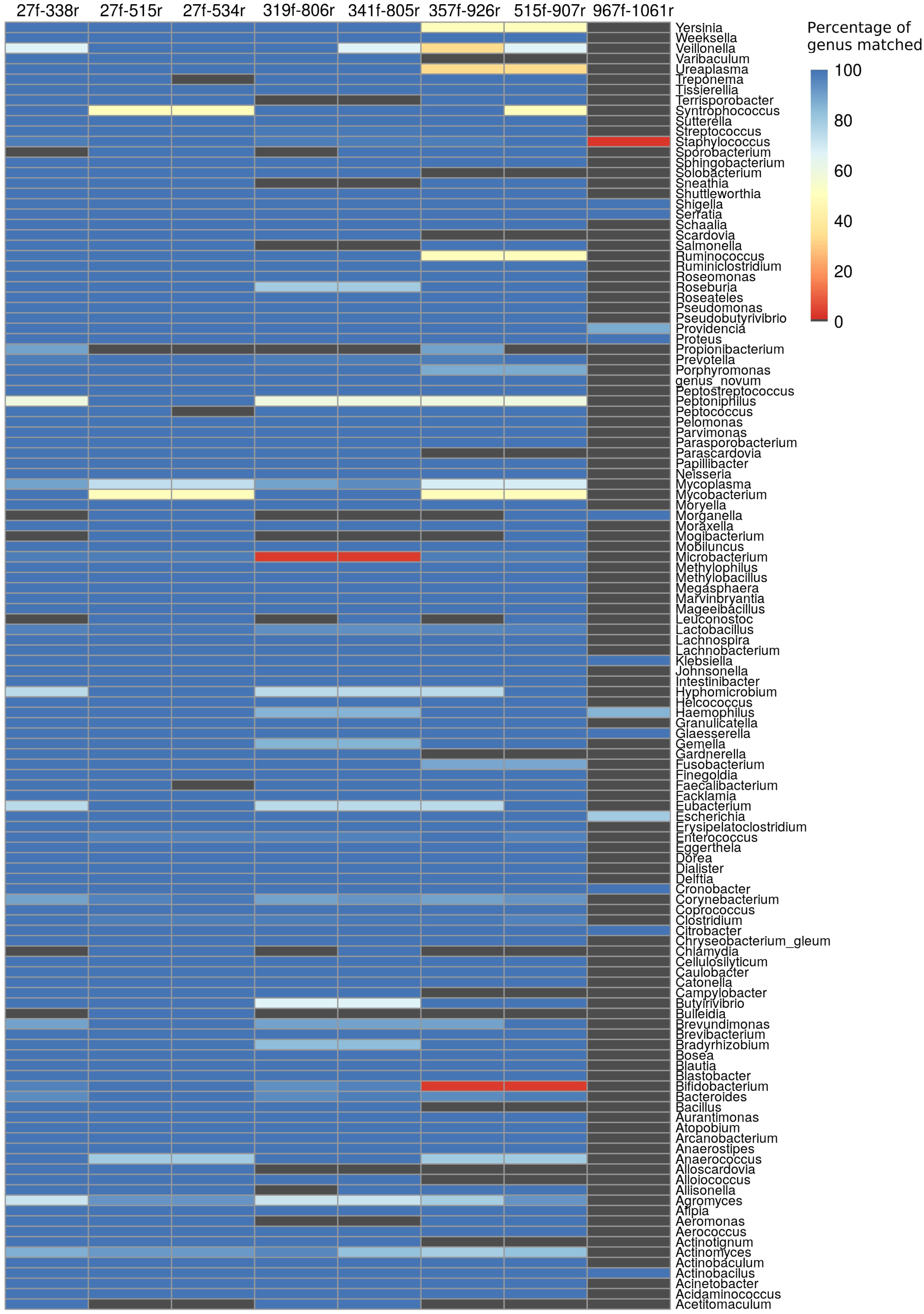
Most previously published work uses 16S primers with good coverage, but a few genera remain a problem, such as *Chlamydia* and *Sneathia*. This heatmap depicts the percentage coverage of each genus (rows) by each primer pair (columns).

### Taxonomic annotation strategies

Even when provided with a primer pair that is potentially informative, researchers must use appropriate bioinformatic pipelines to retrieve this information. At this step, we assume we have perfect error correction capability and do not attempt to simulate PCR and sequencing errors. For long amplicons, where merging of forward and reverse reads might not be possible, we present results for both merged and unmerged reads. **Fig 2a-b** presents the taxonomic accuracy for each primer pair and taxonomic annotation strategy for the full set of vaginal taxa. V1-V2 and V1-V3 perform better for the vaginal microbiome than other regions, provided that they are merged, since processing reads separately entails a loss in precision and accuracy as large as a switch to a different region. Species-level accuracy is particularly critical for genus *Lactobacillus*, since eg *L. iners* is associated to a very different outcome for the host subject than *L. crispatus* (5, 23). **Fig. 2c-d** depicts barplots of taxonomic accuracy for the 114 *Lactobacillus* species included in our study. The trends observed are very similar to the ones observed for the full vaginal database.

**Figure 2.**
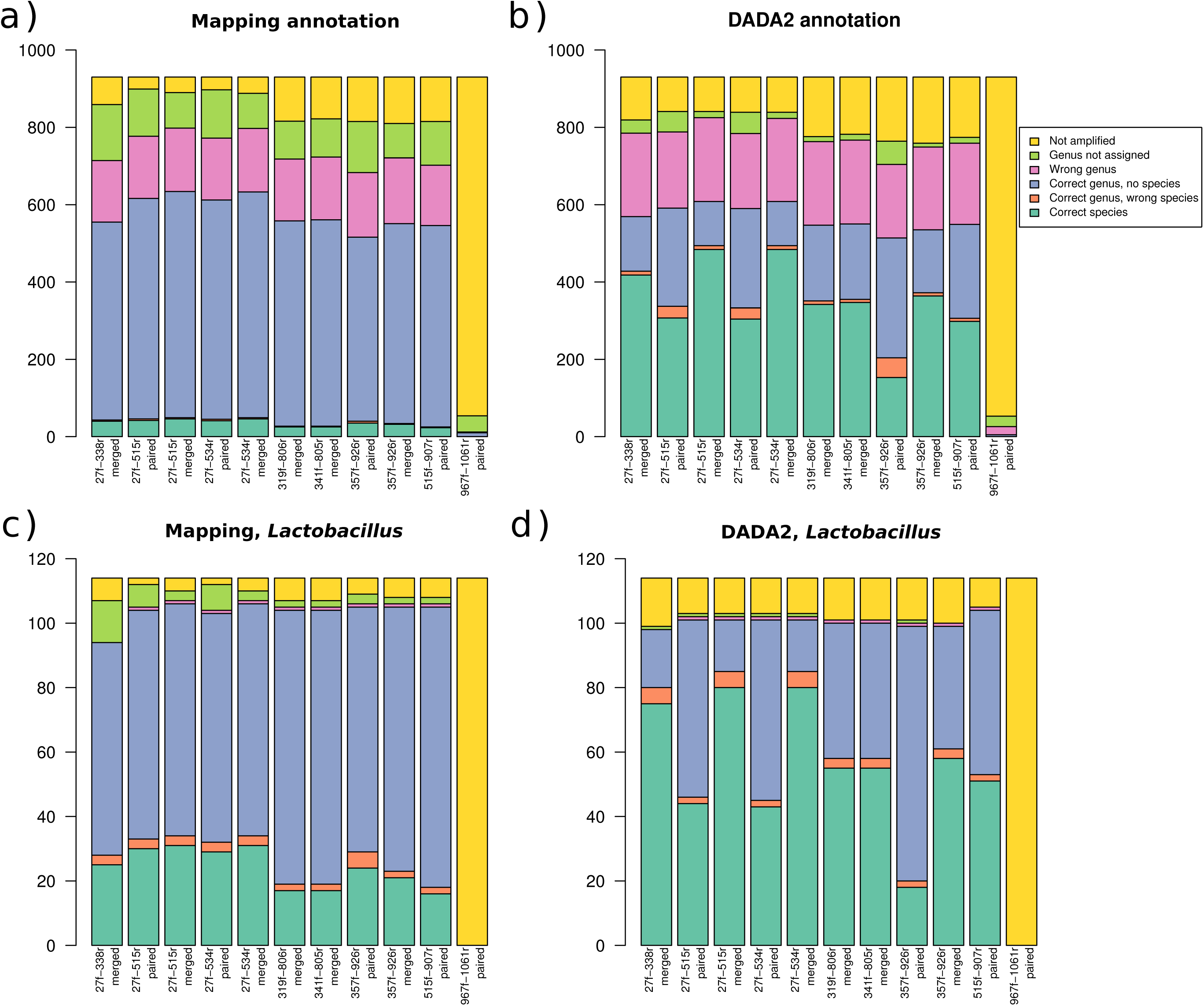
DADA2 taxonomic annotation gives higher taxonomic resolution than mapping to a comparable database, both in general and for *Lactobacillus* in particular. The entire OptiVag database was extracted *in silico* with each of the candidate primer sets, without errors. **(a)** The complete database, annotated by mapping **(b)** The complete database, annotated with DADA2’s algorithm **(c)** Same as (b), but focusing only on *Lacobacillus* **(d)** Same as (a), but focusing only on *Lacobacillus.*

### Amplicon sequencing

To assess the accuracy of these algorithms, 8 pools of vaginal swabs (coming from 4 consecutive days of sampling from a single individual each) were amplified using either the V1-V3 or the V3-V4 region. For the V3-V4 region, two experimental approaches were compared, using either a single PCR reaction (which both amplifies this region and barcodes it), or two consecutive PCR reactions (one for amplification and one for barcoding). The one-step PCR approach is more cost effective, since a single cleaning step is necessary, and minimizes the risk of cross-contamination between wells, since at no point are there samples amplified but not barcoded. However, the long PCR primers can be challenging to obtain, and reaction conditions are also more delicate. Here, both approaches performed very similarly (**fig 3a**), but in some replicates there is a difference in richness (**fig 3b**). This means that either the 2-step approach is producing more artefacts, or that the 1-step approach didn’t capture the full richness of the sample due to worse PCR performance for the long primers. Since the triplicates for the two-step approach yielded more similar results, the latter is the most likely explanation. However, it is also worth considering that while the negative extraction controls for the 1-step approach yielded a total of 3 16S reads post-QC, the 2-step approach had 2,044 reads, highlighting the risk of working with amplified but not barcoded molecules, specially in a high throughput setting.

**Figure 3.**
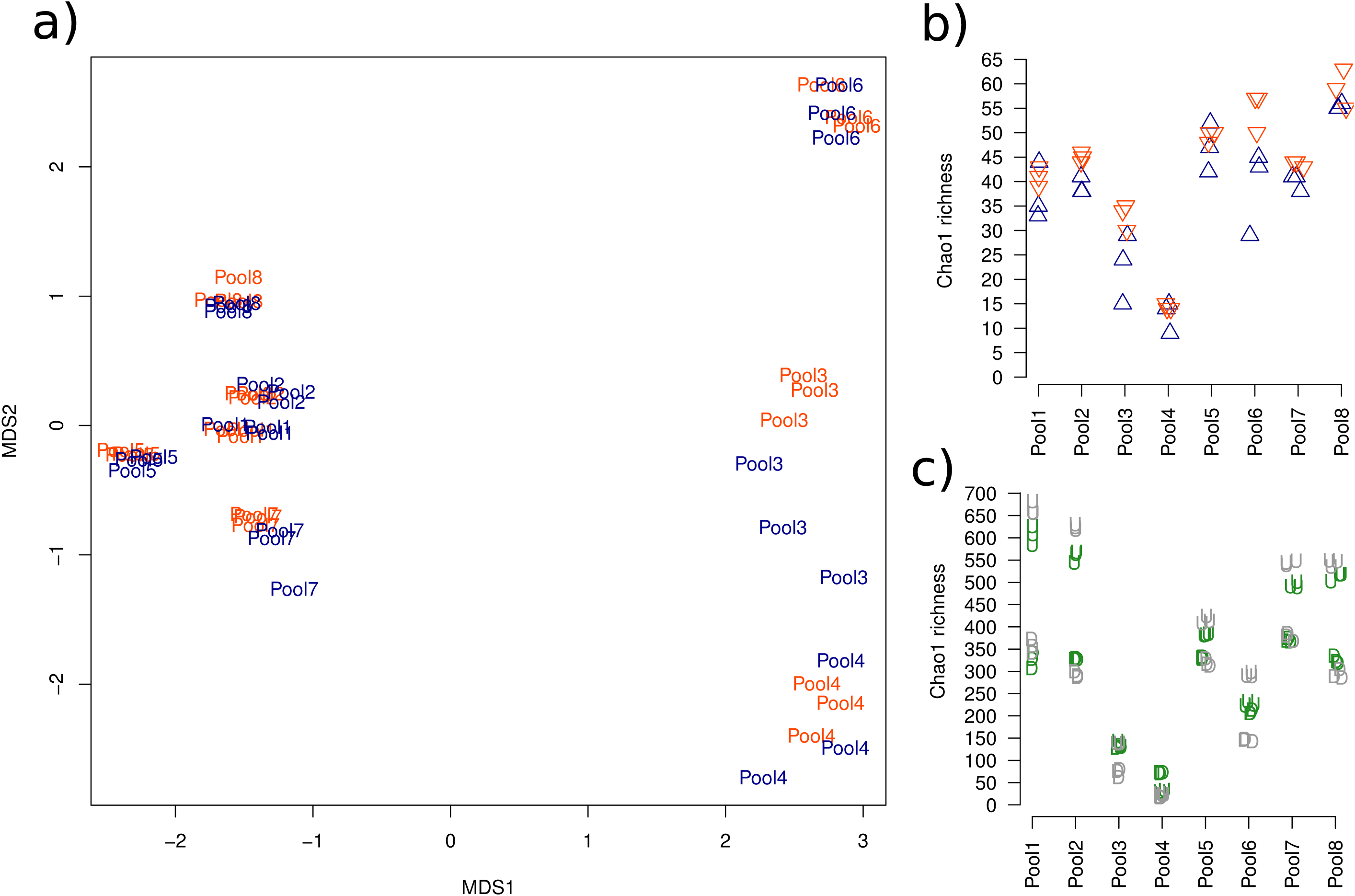
Running PCR in a single step or in two nested reactions has little effect on the results. The DADA2 pipeline yields a more reliable richness estimate than the Unoise annotation when the error rate is high. Orange: V3-V4, 2-step PCR. Blue: V3-V4, 1-step PCR. Green: V1-V3:515r. Gray: V1-V3:534r. **(a)** NMDS of the 8 pools, processed in triplicate with V3-V4 primers, shows good replicability within triplicates and across methods. **(b)** Chao1 richness estimate for each of the same samples in (a). The 1-step PCR approach generally yields a lower richness estimate, but has slightly less replicability within triplicates. **(c)** Chao1 richness of each of the 8 pools amplified in triplicates with V1-V3:515r (green) or V1-V3:534r (gray), and error corrected with DADA2 (D) or Unoise (U).

The V1-V3 amplicons are too long for current paired-end 300 bp approaches to accurately bridge the space between reads, so read concatenation must be used rather than read merging. Failing to merge decreases the accuracy of this middle region, which is generally already low due to the failing accuracy of sequencing along the read length. To achieve species-level resolution, it is crucial to use an error-correction strategy rather than a clustering one. Here, we have compared DADA2 (29) and Unoise3 (30). DADA2 is optimized to correct sequencing errors, and will not eliminate PCR errors, so this algorithm is only recommended in combination with a high fidelity polymerase to avoid large numbers of false positives. Unoise3 eliminates both amplification and sequencing errors, but also presents a higher risk of excluding rare but correct sequences, which generally makes it a more conservative approach (39). However, in the case of non-merged reads, DADA2 seems to be more parsimonious, yielding a lower amount of ASV per sample **(fig 3c)**.

The taxonomic composition of each sample, analysed with the best possible set-up for each primer set, is extremely comparable, but primer pair V1-V3:534r yielded slightly worse taxonomic resolution **(fig 4a)**. Compared to qPCR, all three approaches are extremely accurate **(fig 4b**).

**Figure 4.**
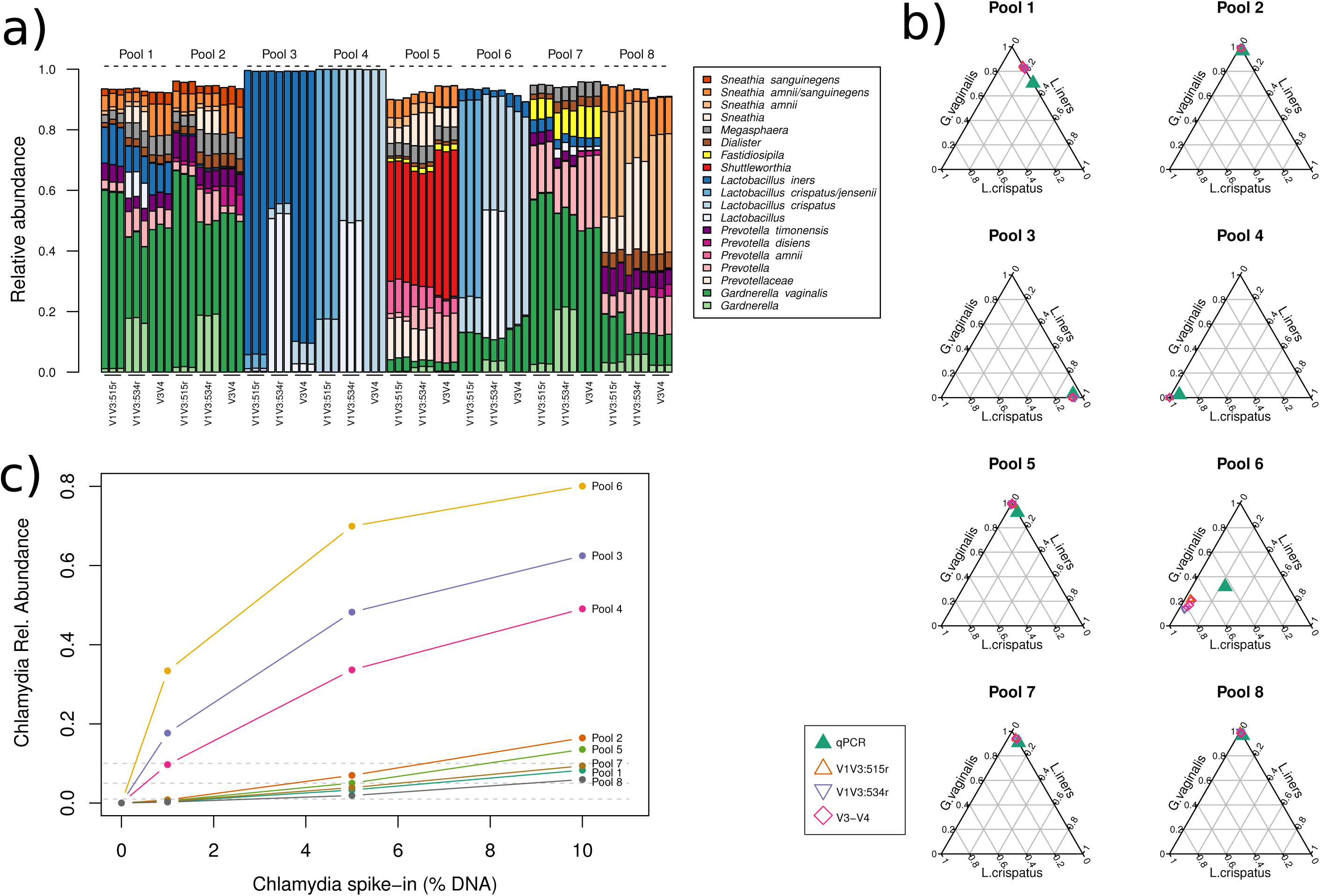
The taxonomic accuracy of each of the 16S primer sets is good, but V1-V3:534r yields more shallow annotations. It can, however, reliably detect *Chlamydia trachomatis* spike-in DNA. **(a)** Taxonomy barplots for each of the pools, processed with 2-step PCR with each of the primer sets in triplicate. **(b)** Same samples as (a), compared to qPCR results for *Lactobacillus iners, Lactobacillus crispatus* and *Gardnerella vaginalis*. For each sample, the sum of these three taxa was normalized to 1, to make them comparable to the qPCR results in the triaxis plot. **(c)** Percentage of *C. trachomatis* detected in each sample as a function of the DNA spike-in. Differences in human DNA content affect the observed bacterial counts, and for the three samples with highest DNA content (Pools 3, 4 and 6), the assay quickly becomes saturated. Dashed grey lines mark 1%, 5% and 10%, which were the proportions used for the spike-in experiment.

Despite its somewhat lower taxonomic resolution with the read lengths obtained, primer pair V1-V3:534r is the only one expected to amplify and detect *Chlamydia trachomatis.* To confirm this, a spike-in experiment was conducted **(fig 4c**). The varying amount of human DNA initally found in each sample means that a spike-in of 5% of total DNA may correspond to >50% of bacterial DNA, making this analysis harder to interpret, but in general there is a good correlation between spiked in and observed *C. trachomatis*.

### *In silico* removal of human DNA from metagenomic data

An alternative to PCR amplification is performing full shotgun metagenomic sequencing of samples. The first challenge for processing metagenomic reads derived from vaginal swabs is the large amount of human DNA in these samples. In our pools, 86-98% of the reads could be mapped to the human genome. While human DNA depletion can be performed *in vitro* prior to sequencing (40, 41), this depends on the storage condition of the samples and was not evaluated in this work. Instead, we focused on *in silico* removal of human reads.

Removal of reads of human origin is a conceptually straightforward process consisting of mapping reads to a reference genome, but two important factors can affect the outcome: the mapping algorithm and the masking applied to the reference genome to hide regions exhibiting homology to Bacteria and Fungi. Strict mapping is time- and memory-intensive. A looser mapping is less resource intensive, but might also remove more bacterial reads or retain more human reads, depending on how strictly the reference is masked. Many mappers provide a pre-formatted human genome reference for host removal. Here we tested three of them: BMTagger, BBmap, and Kraken2, the latter in both “quick” and standard modes. The percentage of human DNA left in each sample after human DNA removal with the different techniques is depicted in **figure 5a**. The percentage of bacterial DNA kept, from the initial bacterial pool, is shown in **figure 5b**. These two quality scores are combined in **figure 5c**, where the optimal method would present all samples in the upper left corner. BBMap and Bowtie2 retained the most human DNA, but also the most bacterial reads. Conversely, BMTagger and Kraken2 removed the most human reads, at the expense of also decreasing the microbial pool. Interestingly, all tools followed the same general trends, removing more bacterial reads and also retaining more human reads in samples with an originally very high (>95%) human DNA content. Of notice, these three samples (Pools 3, 4 and 6) also have the highest *Lactobacillus* counts. Based on the results above, Kraken2 in quick mode was chosen for downstream analysis.

**Figure 5.**
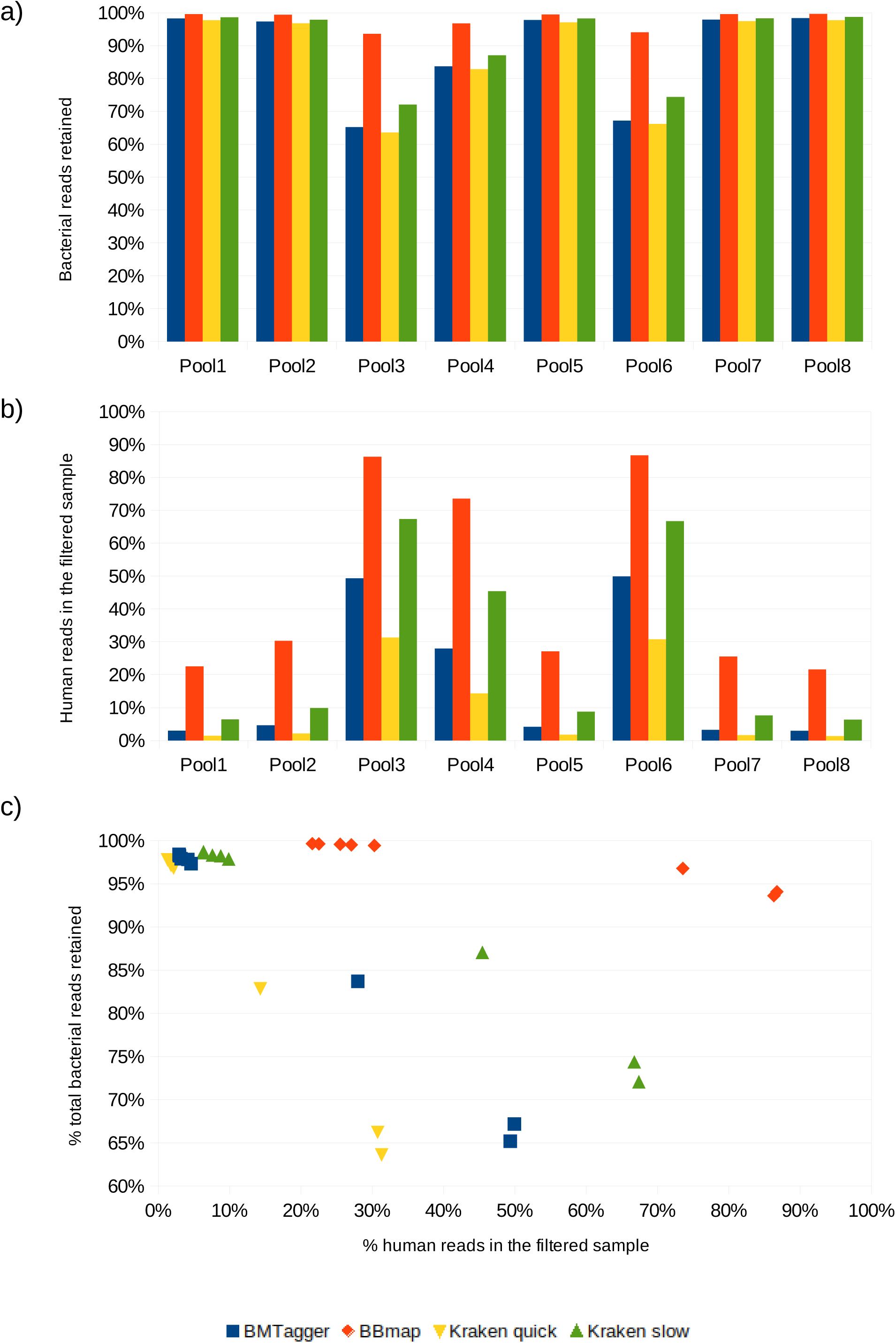
The human DNA content in the 8 pools varied from 86-98% of the total DNA content. Different DNA removal methods leave varying amounts of human DNA in the filtered sample, but also retain varying amounts of the original microbial pool. Kraken2 and BMTagger remove most human DNA, but also the most microbial reads. **(a)** Percentage of human reads in each sample before and after each human removal strategy. **(b)** Percentage of the original pool of microbial reads kept in each sample after each human removal strategy. **(c)** The data from (a) and (b) is plotted against each other for an easier overview of each tool.

### Taxonomic annotation of metagenomic data

Five approaches were assessed for taxonomic assignment on these data: a general marker-gene based approach (MetaPhlAn2), a marker-gene based approach built from a curated set of vaginal bacteria (VIRGO), a k-mer based approach with a broad taxonomic database (Kraken2, see methods for details), a k-mer based approach with a vaginal-only database (Kraken2), and a novel pre-filtering and alignment tool (Metalign). The taxonomic profile inferred by each method for each pool is depicted in **figure 6a**. Metalign stands out in identifying *Chlamydia trachomatis* in almost every pool, as well as a higher frequency of detection of *Veillonela spp.* and *Prevotella spp.* The standard Kraken2 database failed to identify *L. iners*, despite this species being present in the database. OptiVag, Metaphlan and VIRGO tend to present similar results, with a few notable differences. Firstly, the clade called BVAB3 in VIRGO takes its current name *Mageeibacillus indolicus* in the other two references. Metaphlan fails to identify BVAB1, perhaps because this genome is still not in NCBI’s RefSeq database. OptiVag is alone in identifying significant amounts of *Peptoniphilus* in three of the *Gardnerella*-dominated samples. This clade has been previously identified in women with bacterial vaginosis (42), but is generally not considered a key taxon for this condition. Finally, VIRGO stands out in not identifying any *Sneathia*, even in samples where all other methods are in agreement.

**Figure 6.**
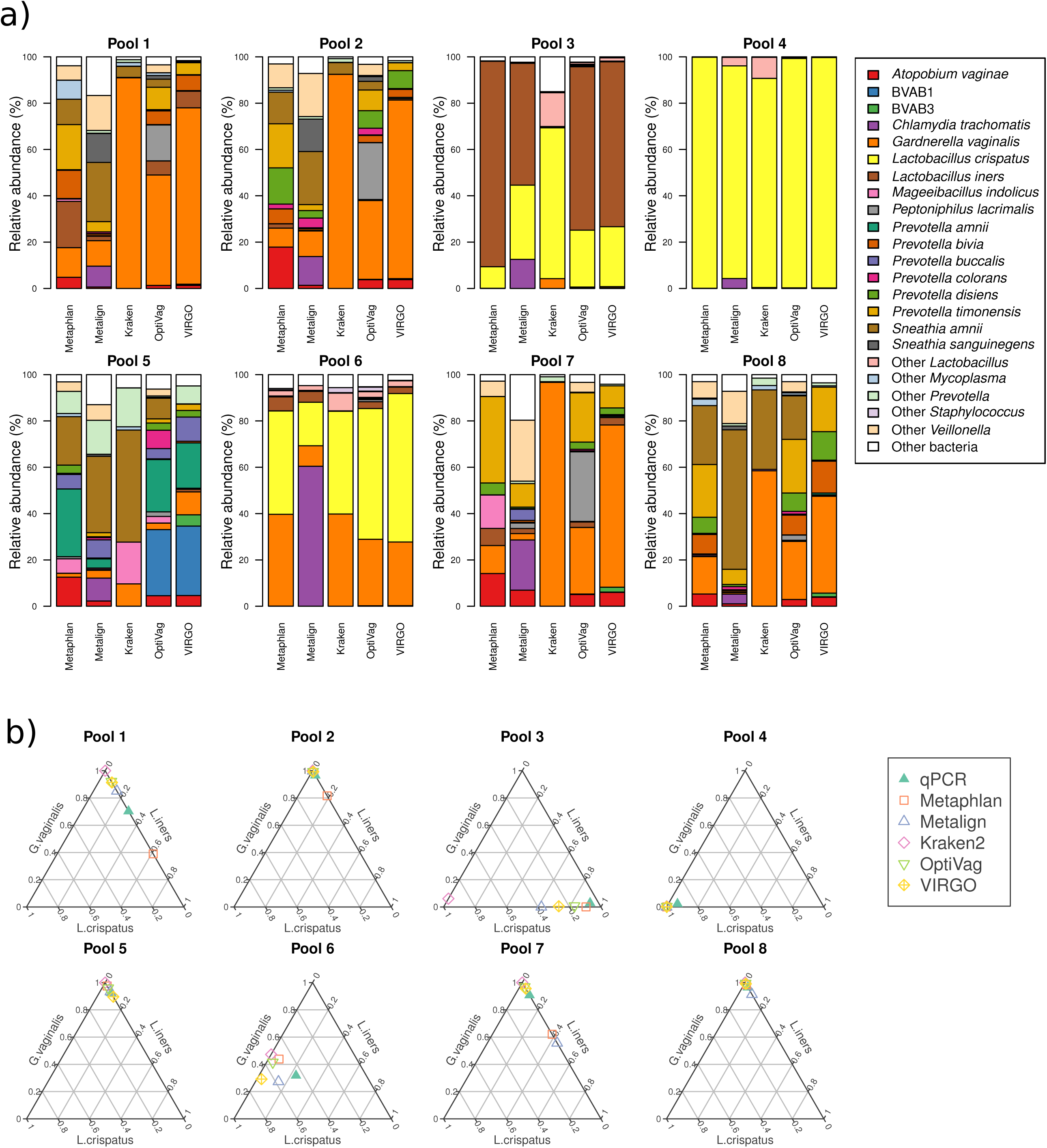
Assigning taxonomy to shotgun metagenomic reads with various tools yields somewhat different community profiles. **(a)** Taxonomy for each pool assigned with Metaphlan, Metalign, Kraken2 to its complete microbial database, Kraken2 to the OptiVag database or VIRGO. **(b)** Same samples as (a), compared to qPCR results for *Lactobacillus iners, Lactobacillus crispatus* and *Gardnerella vaginalis*. For each sample, the sum of these three taxa was normalized to 1, to make them comparable to the qPCR results in the triaxis plot.

When comparing to the qPCR results, none of the shotgun methods were as accurate as the PCR-based methods **(fig 6b;** contrast to fig. 4b). Still, when considering each of the pools, VIRGO and OptiVag performed better than the other methods (fig. 4c). It is possible that assessing taxonomy after assembly would yield more accurate results (43), but this was not possible with the current sampling depth. Still, this could be a valid alternative for samples sequenced more deeply, or for a different experimental design, eg a time-series from the same woman, which would enable co-assembly across closely related samples.

## Conclusions

None of the methods’ assessed here are superior in all aspects. In regards to amplicons, V3-V4 yielded the most plausible alpha-diversity estimates and has very good taxonomic coverage. However, much of the existing literature is based on region V1-V3 (14–16). The major drawback of 16S amplicons is their failure to detect eukaryotic taxa such as *Candida spp.* and *Trichomonas vaginalis*. An ITS-based amplicon approach could selectively amplify fungi without amplifying human DNA (44), but it would miss the pathogenic Parabasalid *T. vaginalis.* Therefore, no simple combination of one or two primer sets can accurately profile all relevant taxa in the human vaginal environment.

To overcome such limitations, shotgun metagenomic sequencing presents an interesting alternative, since it is not *a priori* bound by phylogeny. Its cost, which used to be prohibitive, is now low enough to compete with a multiprimer PCR-based approach. In addition to taxonomic classification, shotgun data also allows researchers to assess the functional gene content of a sample and, given enough sequencing depth, assemble draft genomes of strains of interest.

The main practical obstacle to a broader application of shotgun metagenomics in the field of obstetrics and gynecology is the high amount of human DNA in vaginal swabs, but this can potentially be bypassed, either with molecular biology techniques or a combination of deep sequencing and *in silico* human DNA removal. The bioinformatics skillset and computational requirements necessary to handle this type of data is also significantly larger than what is needed for marker gene (16S) analyses.

Here we have presented a thorough comparison of multiple methods available for the survey of the vaginal microbiota. Since none of them are universally optimal, it is still up to each research center to select the appropriate method for their specific research question. While this will necessarily limit comparability between studies, acknowledging the strengths and weaknesses of each method is already a large improvement to the current state of the field.

## Author contribution

LWH - wrote code, performed *in silico* experiments, planned *in vitro* experiments, analysed the sequencing data, wrote the manuscript

MP, YZ, MS - planned and performed *in vitro* experiments, wrote the manuscript VK - wrote code, performed *in silico* experiments

FB, ISK, MH – planned experiments, wrote the manuscript EF, LE – wrote the manuscript, supervised students

MCK, ZB and HSN – Obtained ethics and data protection approval, planned and organized the study cohort, included participants and secured informed consent and collected samples All authors have read and approved the final manuscript

## Acknowledgments

We thank Pia Angelidou from the Centre for Translational Microbiome Research for her efforts in DNA extraction.

---

| Region | Primer_fw | Primer_fw | Primer_rev | Primer_rev | Reference |
| --- | --- | --- | --- | --- | --- |
| V1-V2 | 27f | AGAGTTTG | 338r | GCTGCCTC | Ravel, 2011; Brotman, 2014 |
| V1-V3 | 27f | AGAGTTTG | 534r | ATTACCGC | The |
| V1-V3 | 27f-pool | 4xAGAGT | 515r | TTACCGCG | Frank, 2008; Hugerth, 2014 |
| V1-V3 | 27f-pool | 4xAGAGT | 533r | ATTACCGC | Frank, 2008 |
| V3-V4 | 319f | ACTCCTRC | 806r | GGACTACH | Elovitz, 2019 |
| V3-V4 | 341f | CCTACC | 805r | GACTACHV | Hugerth, 2014 |
| V3-V5 | 357f | CCTACC | 926r | CCGTCAAT | Jumpstart |
| V4-V5 | 515f | GTGCCAGC | 907r | CCGTCAAT | Chen, 2017 |
| V6 | 967f | CAACGCGA | 1061r | ACAACACG | MÃandar, 2015 |

| Primer | Counts | Percentage |
| --- | --- | --- |
| 27f " sir | 249 | 26.80% |
| 27f " poc | 297 | 31.90% |
| 319f | 896 | 96.30% |
| 338r | 909 | 97.70% |
| 357f | 892 | 95.90% |
| 515fr | 898 | 96.60% |
| 534r | 896 | 96.30% |
| 806r | 848 | 91.20% |
| 907/926r | 842 | 90.50% |
| 967f | 670 | 72.00% |
| 1061r | 55 | 5.90% |

| Primer | Proportio |
| --- | --- |
| 27f-338r | 26.10% |
| 27f-338r | 95.70% |
| 27pool- | 30.90% |
| 27pool- | 96.60% |
| 27f-534r | 25.60% |
| 27f-534r | 96.30% |
| 27pool- | 25.60% |
| 27pool- | 96.30% |
| 319f-806r | 87.80% |
| 341f-805r | 88.50% |
| 357f-926r | 87.60% |
| 515f-907r | 88.10% |
| 967f-1061r | 67.30% |

